# Assessing the impact of artificial night lighting regulations designed to protect astronomical observatories on seabirds and bats

**DOI:** 10.64898/2026.04.16.718868

**Authors:** Cristina de Tena, Beneharo Rodríguez, David Garcia, Juan Federico de la Paz, Airam Rodríguez

**Affiliations:** Departamento de Ecología Evolutiva, Museo Nacional de Ciencias Naturales (MNCN-CSIC), C/ José Gutiérrez Abascal 2, 28006, Madrid, Spain; Canary Islands’ Ornithology and Natural History Group (GOHNIC), C/ La Malecita s/n, Buenavista del Norte 38480, Tenerife, Canary Islands, Spain; Islands Biodiversity Research Initiative (IRBI), Plaça Nova 30, Alaior, 07330, Menorca, Balearic Islands, Spain; Oficina Técnica para la Protección de la Calidad del Cielo, Instituto de Astrofísica de Canarias, C/ Vía Láctea s/n, 38205, Tenerife, Canary Islands, Spain

**Keywords:** Light pollution, attraction, disorientation, light intensity, wavelength, behaviour, marine birds, bats

## Abstract

Artificial light at night is a rapidly increasing driver of global change, affecting both astronomical observations and biodiversity. Regulations such as the Canary Islands’ “Sky Law” were designed to protect astronomical observations by controlling light intensity and spectral composition, yet their ecological effectiveness remains largely untested. Here, we experimentally assessed whether lighting conditions permitted under this law influence the behaviour of two sensitive nocturnal taxa: seabirds and bats. Field experiments were conducted in Tenerife, Canary Islands, using controlled lighting treatments that varied in intensity (low vs. high) and spectrum (PC amber ~1800K vs. white ~2700K), including a no-light control. We monitored the behaviour of breeding adult Cory’s shearwaters (*Calonectris borealis*) using GPS tracking and passive acoustic recording, and quantified bat activity through ultrasonic detectors. Behavioural responses included flight characteristics, colony attendance, vocal activity in shearwaters, and species-specific movement and feeding activity in bats. Generalised linear mixed models were used to evaluate treatment effects while accounting for environmental covariates. Across 211 shearwater flights and extensive acoustic datasets, we found no consistent or significant effects of light treatments on seabird flight behaviour, vocal activity, or bat movement and feeding activity. Instead, environmental variables such as moonlight, seasonality, and interannual variation were stronger predictors of behavioural responses. These results suggest that lighting conditions currently permitted under the Sky Law may have limited ecological impact on the studied taxa under the conditions tested. Further research in less disturbed environments and with broader spectral contrasts is needed to better assess the ecological implications of astronomically motivated lighting regulations.

## 1. INTRODUCTION

Light pollution hinders observation of the night sky and has been one of the biggest challenges facing the field of astronomy for a long time [1,2]. Unsurprisingly, astronomers were who first campaigned for laws to regulate alterations to natural night-time light levels caused by artificial light [3]. Later, conservationists drew attention to the natural loss of the sky, and the medical community pointed to the control of light levels to respect natural biological rhythms [4]. Despite these efforts, light pollution is a global phenomenon increasing at an alarming rate in recent decades [5] with sky brightness growing by between 2.2% and 9.6% each year [6,7]. This makes light pollution one of the fastest-growing drivers of global change [8].

Most organisms have evolved under a relatively stable natural cycle of light and darkness [9]. Thus, artificial light disrupts the way in which organisms perceive and respond to the natural light, a key driver in the evolution of biological rhythms [8]. This disruption leads to a wide range of ecological impacts with cascading effects on the functioning of ecosystems [9–11]. Thus, light pollution has become a significant threat to the conservation of biodiversity, affecting not only nocturnal species, but also diurnal ones and in all ecosystem realms [12–14]. Among the many effects, mass mortality events, such as those affecting flying insects, sea turtles and seabirds, are among the most dangerous in the short term [10].

As awareness of the ecological impacts of light pollution grows, understanding how light properties (i.e. intensity, spectrum, timing, and spatial distribution) affect ecosystem functioning becomes increasingly important [15]. Among these properties, intensity and spectrum are particularly influential yet remain underexplored. Investigating how variations in these properties influence biodiversity offers a valuable opportunity to assess and improve such policies, drawing on mitigation strategies [16].

The national law for the protection of the astronomical quality of the observatories of the *Instituto de Astrofísica de Canarias* (IAC), hereafter Sky Law, is a pioneering piece of legislation not only in Canary Islands and Spain, but worldwide in the field of light pollution regulation (Law 31/1988). This law regulates light characteristics in terms of intensity and spectra emitted in La Palma and Tenerife Islands, ensuring the protection of astronomical observations conducted from the IAC observatories. Like most laws promoted by the astronomy community, this legislation was developed for safeguarding dark skies for observational purposes. However, protecting the night sky should not be limited to astronomical interests; broader environmental, ecological and social dimensions should also be considered [17]. Proactive management must be integrated into both conservation efforts and lighting design. This calls for a transdisciplinary approach that connects scientific understanding with policy development and community involvement [11,15].

Canary Islands is the richest region in Spain and among the richest in Europe in terms of biodiversity, endemism, and rarity [18]. Therefore, it is particularly important to experimentally evaluate whether the restrictions set out in the astronomic-based Sky Law of the Canary Islands are effective in reducing ecological impacts of light pollution on wildlife. To this end, we focus our research on two animal groups that have been widely studied and are particularly sensitive to light pollution: nocturnal procellariform seabirds and insectivorous bats. Our aim is to investigate the impact of light pollution on their activity and behaviour, and ultimately improve their conservation.

For seabirds, particularly nocturnal Procellariiformes, light pollution poses a significant threat recognized since a long time ago [19]. Fledglings are particularly vulnerable during their first flights when they are attracted to and disoriented by artificial lights, often landing in areas that are unsuitable for them with fatal consequences [20]. Although adults are generally less affected by artificial light, they suffer mass mortality events, similar to those of fledglings. One of the most surprising cases is that of thousands of auklets colliding with a fishing boat and causing it to list so severely that it nearly capsized [21] or that of 131 endangered seabirds grounded at a single Air Force Station on Kauai, Hawaii [22]. Artificial lighting near colonies can also alter nocturnal attendance behaviour, delay landings and potentially affecting nourishing in adult fairy prions [23], and Scopoli’s and Yelkouan shearwaters [24,25].

For bats, artificial night lighting have multiple impacts, exhibiting divergent behavioural responses to light [26,27]. Some bat species are especially sensitive to disturbance from artificial light, as their behaviour and foraging activity are affected by moonlight whose light levels are usually overridden by artificial light levels [28]. The light spilling over from urban areas into potential bat habitats can reduce their suitability and disrupt the natural behaviour of the resident species [29]. Excessive illumination can fragment critical habitats by breaking up essential routes between roosting and feeding sites [30]. This fragmentation can compromise key ecological functions such as foraging efficiency, breeding success, and long-term population stability [31,32]. By contrast, other bat species increase their foraging opportunities near light sources, taking advantage of the concentration of light-attracted insects [33,34].

In our study, we experimentally tested the effects of light intensity and spectral composition on (1) the behaviour (flight characteristics and vocal behaviour) of breeding adult Cory’s shearwaters (*Calonectris borealis*) during inward flights to attend the colony using GPS tracking and passive acoustic monitoring; and (2) the species-specific movement and feeding behaviour of the bat community assessed through the ultrasonic pulses recorded by acoustic recorders. We compared combinations of high and low intensity light, with PC amber (~ 1800K) and white LED light (2700K), i.e., the most permissive value of Correlated Color Temperature (CCT) of the restrictions of the Sky Law (Law 31/1988).

Given that light emitted at wavelengths below 500 nm (blue) is particularly disruptive to many taxa [35–37], we hypothesised that seabird flights would be characterised by reduced speed, increased turning angles and tortuosity, extended duration, delays in the time of arrival to the nest, further distance and higher altitude to experimental light, and fewer visits to the colony in conditions of higher intensity and CCT light, i.e. with the presence of more intense-white light. Furthermore, we expected a reduction in vocalisations due to the increased exposure to experimental light levels in the same line with the typical reduction of vocal activity at colonies of nocturnal procellariiforms during nights with high moonlight intensity [38]. For bats, we predicted that light-attracted flying insects would trigger a response from certain insectivorous bat species, particularly light-tolerant or opportunistic genera, such as *Pipistrellus* and *Nyctalus*, increasing rates of echolocation pulses and feeding buzzes under high-intensity and short-wavelength light. Conversely, light-averse bat species, including more light-sensitive genera would be less present in the experiment-illuminated area or would reduce their vocal activity compared to darker conditions (control or low-intensity and long-wavelength light treatments).

## 2. METHODS

### 2.1. Study area and model species

Our field experiment was conducted from July to August of 2023 and 2024 at Reserva Natural Especial Barranco del Infierno, located at south-western part of Tenerife, the largest island of the Canary Islands. Tenerife Island holds approximately 1 million residents and also attracts 7 millions of tourists each year that occupy mostly the southern part [39]. Barranco del Infierno is a Natura 2000 protected area (*Reserva Natural Especial*) with regulated access to the public (https://barrancodelinfierno.es).

Cory’s shearwater (*Calonectris borealis*; Cory, 1881) is a medium sized procellariform seabird (body mass: 605-1,060 g, and wingspan: 113-124 cm) that nests on North Atlantic Ocean islands, mainly in Azores, Madeira, Selvagens, and the Canary Islands [40]. It is a long-lived, monogamous seabird that returns each year to the same underground nest to reunite with their mate and lays a single egg per breeding season [40]. This species is strongly affected by artificial light at night. The impact causes high mortality among fledglings, and, under certain conditions (such as rain or fog), adults may also be affected [41]. 6% and 14% of the fledglings annually produced in the archipelagos of Azores and Canary Islands are grounded by artificial light [42,43].

Bats play an essential ecological role as opportunistic insectivorous species that feed on insects, contributing to the natural control of insect populations [44,45]. Bats adapt to both natural and human-modified environments and exhibit primarily nocturnal activity. In Tenerife, seven bat species have been recorded, making it the island with the highest bat diversity in the Canary Islands [46]. Although relatively little is still known about the ecology and behaviour of the Canarian bats, five species are present in our study area and consequently susceptible to be subjects in our experimental study. Artificial light has increasingly become a threat to bats worldwide. Although, no specific studies have been conducted on the Canaries, it is expected that Canarian bats respond to light pollution in a similar way to those from overseas, i.e. affecting foraging behaviour, disrupting commuting routes, and increasing predation risk [32].

### 2.2 Experimental LED lights procedure

Experimental lights were installed in a public square in Barranco del Infierno, Adeje (28.126317, −16.723569), oriented toward the ravine depression, where the seabird colony is located and where bats forage (Figure 1 and <u>Figure S1</u>). Once installed, the lights were programmed to show five different treatments varying in wavelength and intensity: no light or control (C), amber light at low intensity (AL), amber light at high intensity (AH), white light at low intensity (WL), and white light at high intensity (WH) (<u>Figure S1</u>). These treatments were applied randomly on alternating nights (only one treatment per night) throughout the study period (see <u>Table S1</u>). Lights were on since sunset to midnight (local time). LED projectors GALILEO 1 (0I24 ASP-7W A.7-3M) and AREAFLOOD PRO S (AFP S 36L70-827 A6 HFX CL1 GY) were used to produce PC amber (~1800K) and white (2700K) light. These luminaires were produced by AEC Illuminazione and Thorn Lighting, respectively, and set to yield 9280 lm and 18560 lm at low and high intensity, respectively.

**Table 1.** Status of the bat species present in the Canary Islands. Conservation status follows the IUCN European Red List, while island distribution and abundance refer to the Canary Islands and Tenerife, respectively. Island codes: P = La Palma; G = La Gomera; T = Tenerife; C = Gran Canaria; F = Fuerteventura; L = Lanzarote; H = El Hierro. ᵘ Suspected presence, pending confirmation. ᵛ Presence reported, pending validation.

| Scientific name | Common name | Conservation status | Island distribution | Abundance status Tenerife | Biogeographic status |
| --- | --- | --- | --- | --- | --- |
| *Pipistrellus kuhlii* | Kuhl's Pipistrelle Bat | Least Concern (LC) | G<sup>u</sup>, T<sup>v</sup>, F | Rare | Palearctic |
| *Pipistrellus maderensis* | Madeiran Pipistrelle Bat | Vulnerable (VU) | P, G, T, H | Abundant | Macaronesia endemic |
| *Hypsugo savii* | Savi's Pipistrelle Bat | Least Concern (LC) | P, G, T, C, H | Abundant | Palearctic |
| *Nyctalus leisleri* | Leisler's Bat | Least Concern (LC) | P, T, H | Abundant | Palearctic |
| *Barbastella barbastellus guanchae* | Canary barbastelle | Vulnerable (VU) | G, T | Scarce | Canary endemic |
| *Plecotus teneriffae* | Canary Long-eared Bat | Critically Endangered (CR) | P, T, H | Common | Canary endemic |
| *Tadarida teniotis* | European Free-tailed Bat | Least Concern (LC) | P, G, T, C | Abundant | Palearctic |

**Figure 1.**
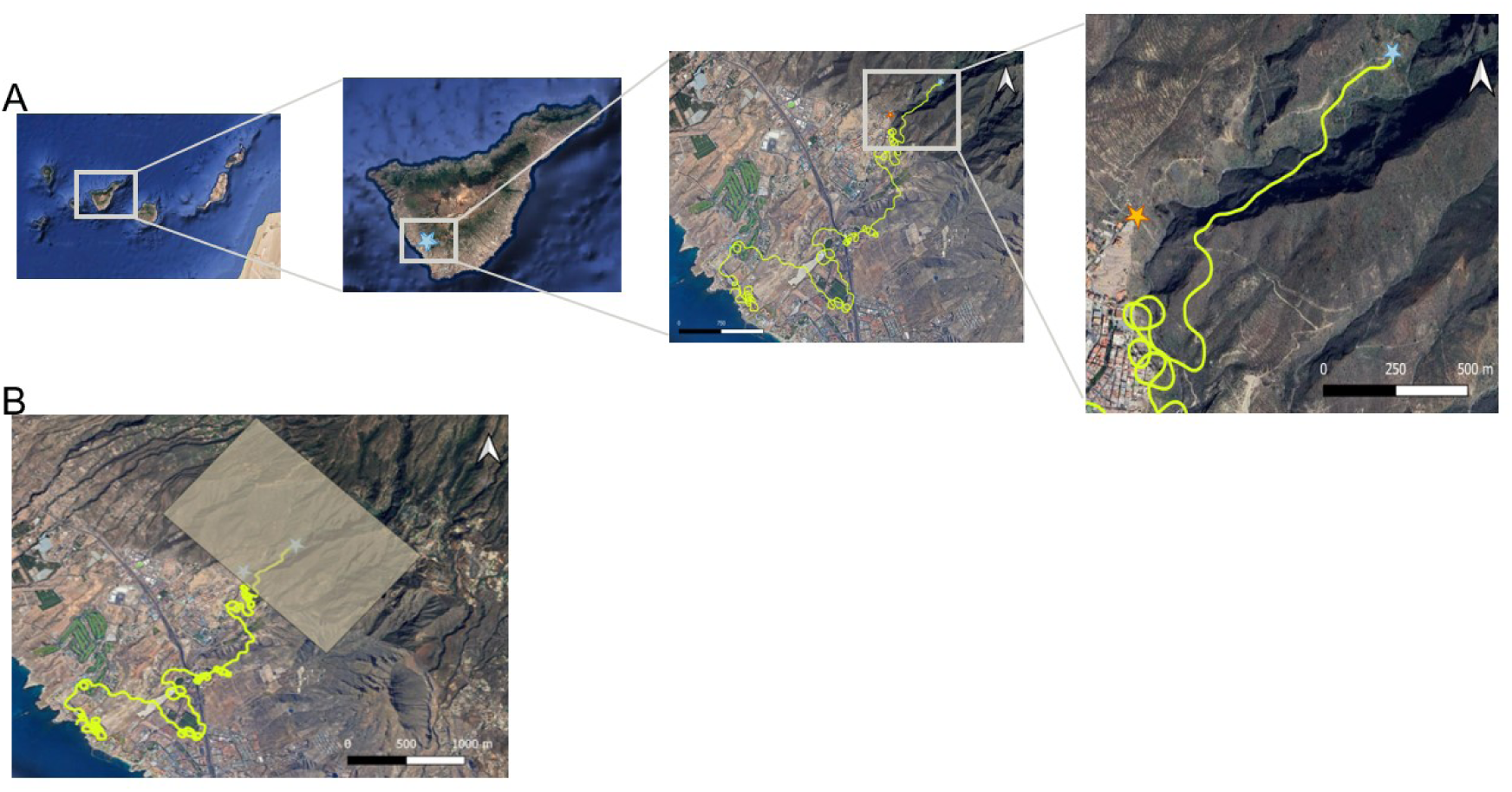
The four upper pictures show the study area (A). The first one indicates the location of Tenerife Island in the Canary Archipelago. The second picture shows the location of the study area (blue star) in the south-west of Tenerife Island. The third shows with a yellow line the GPS-tracked flight of an adult breeder Cory’s shearwater (*Calonectris borealis*) from the sea to its nest at a temporal resolution of 1 fix/second. The fourth picture display the GPS-track in the experimental area. Orange star indicates the location of experimental lights and blue star indicates the nest. The fifth picture (B) shows the area considered to the GPS-track analyses.

### 2.3 GPS tracking data

To assess the effect of the light treatments on the behaviour of adult shearwaters during their inward commuting flights to the colony, we captured 26 breeders (14 in 2023 and 12 in 2024 during the chick rearing periods) by hand in their nests and deployed with remote-download GPS devices (solar powered OrniTrack-9). The GPS devices weight less than 10 grams, which is less than 2% of the body mass of adult breeders, and were tagged to the plumage of the back attached with TESA tape [48]. GPS devices were programmed to record one location per second when the animal is within predefined polygons (experimental study area; see below and Figure 1B), and one location every five minutes when outside these areas (at sea) to safe battery.

As a first step in the data filtering process, the recorded tracks analyses were conducted in QGIS software version 3.10. The pre-processing of these data involved defining a rectangular area (with a total surface area of 10 km^2^) encompassing both the experimental artificial lights and the breeding colony (study area; Figure 1B). Tracks that were interrupted due to GPS malfunctioning or beyond the boundaries of the predefined polygon were excluded, and the remaining points were subsequently classified into distinct daily tracks. QGIS enabled a spatial visualization of the GPS-tracks, facilitating the identification and classification of inward and outward flights. Shearwaters return from their foraging areas to the colony mainly during the first hours of the night (note Cory’s shearwaters only visit their nest at night). Accordingly, light treatments were switched on since sunset until midnight (see below) to coincide with most colony visits. This part-night lighting setting was a requirement of the bioethics committee allowing potential disturbed birds visiting the colony later at night when lights were turnoff (after midnight). Thus, we focussed on inward flights when birds are strongly motivated to reach the nest and feed their chicks.

We obtained eight response variables from each inward flight: speed (m/s), turning angle (turns; deg), duration of the flight (time; secs), tortuosity index, arrival time since sunset (TSS; secs), nearest distance to light (NDL; m), altitude at the NDL (m), and number of visits per night (NV) (Table 2). For speed, turning angle, and tortuosity index, the first and last GPS locations were those when birds entered in a predefined colony polygon (Figure 1B) and arrived at their nests.

**Table 2.**
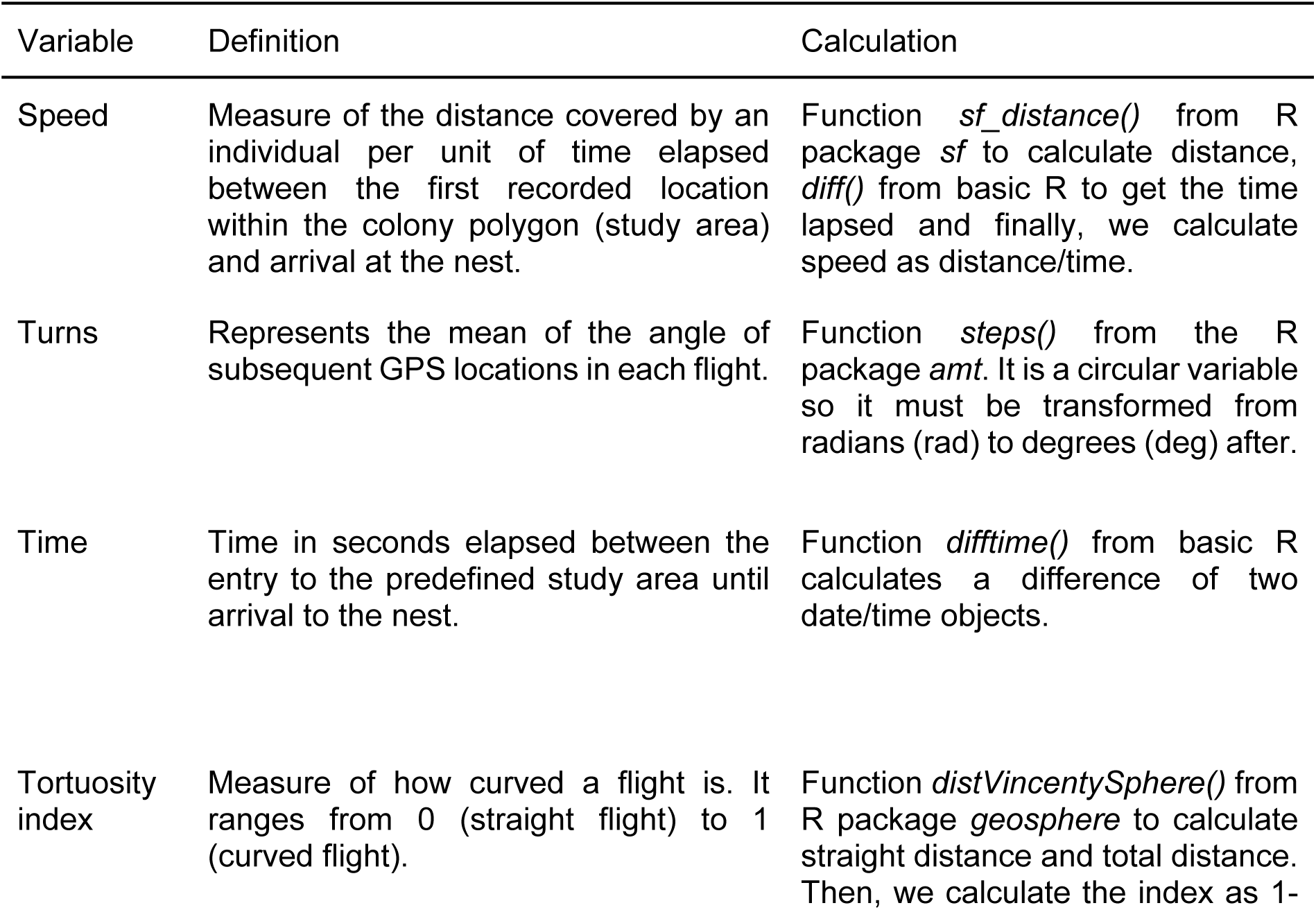
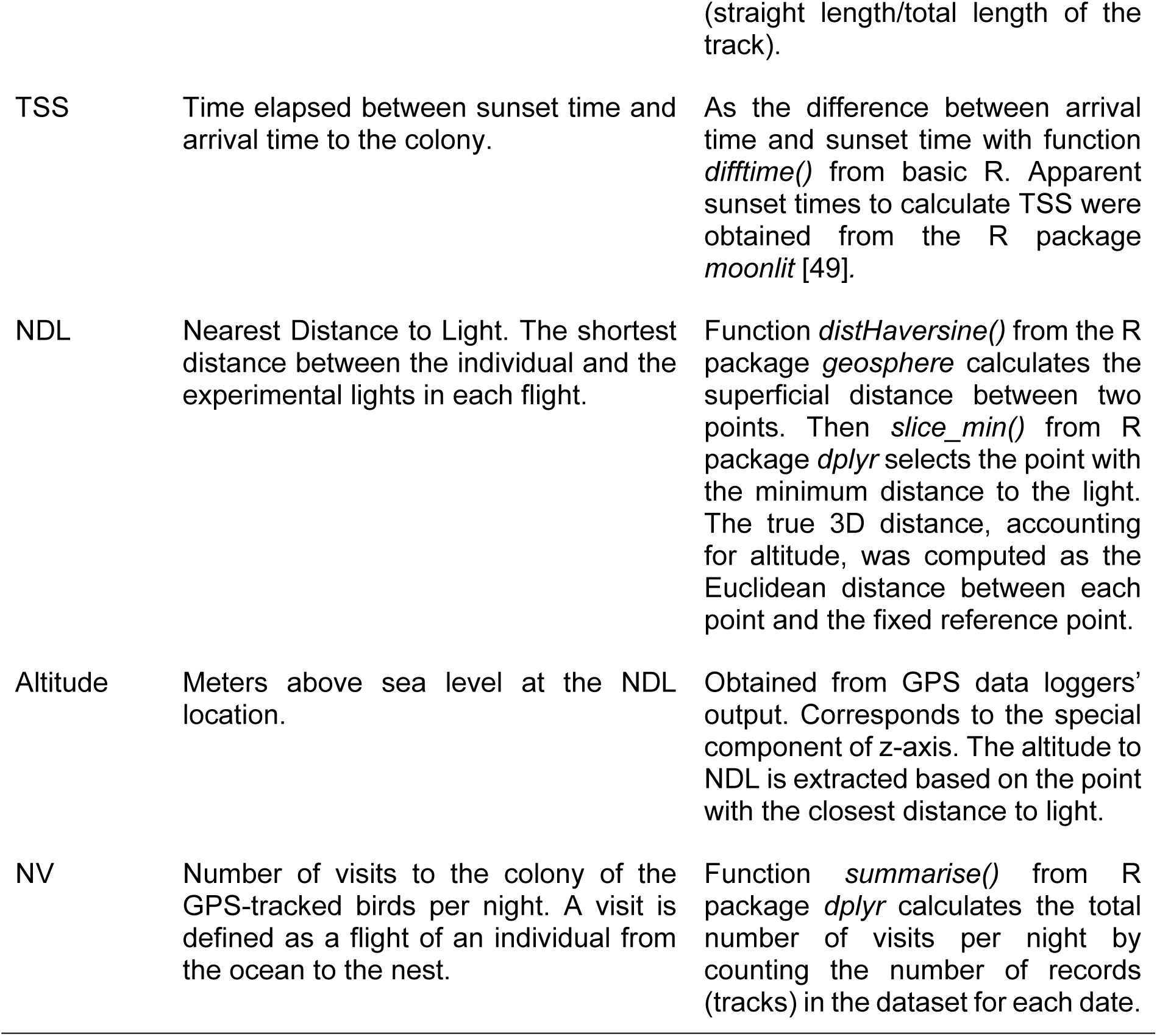
Description of the response variables obtained from the GPS data used for statistical analyses.

### 2.4 Acoustic recording

To monitor the activity of seabird and bat species, two different acoustic recording systems from Wildlife Acoustics (Maynard, USA) were deployed near the light sources. Vocalizations of Cory’s shearwaters were recorded using two autonomous audio recorders (Song Meter Micro) with built-in microphones, while bat ultrasonic echolocation calls were recorded using SM4BAT detectors equipped with external ultrasonic microphones and operating in full-spectrum mode (sampling rate 384 kHz). Recordings were triggered automatically and stored as files of up to 5 s in duration. Bat detectors were configured to capture ultrasonic signals within a frequency range of 8–120 kHz, covering the echolocation calls of the bat species present in the Canary Islands.

Cory’s shearwater vocalizations occur within the audible frequency range, mainly between 2 and 10 kHz. Recordings were conducted at a sampling rate of 48 kHz, and subsequent analyses focused on the 2–8 kHz range, corresponding to the main energy band of Cory’s shearwater vocalizations. The devices remained active throughout the experimental period during which light treatments were applied, recording from sunset to midnight. Recordings were analysed as proxies for three variables (Table 3): Cory’s shearwater vocal activity rate (VAR), bat movement activity, and bat feeding activity.

**Table 3.**
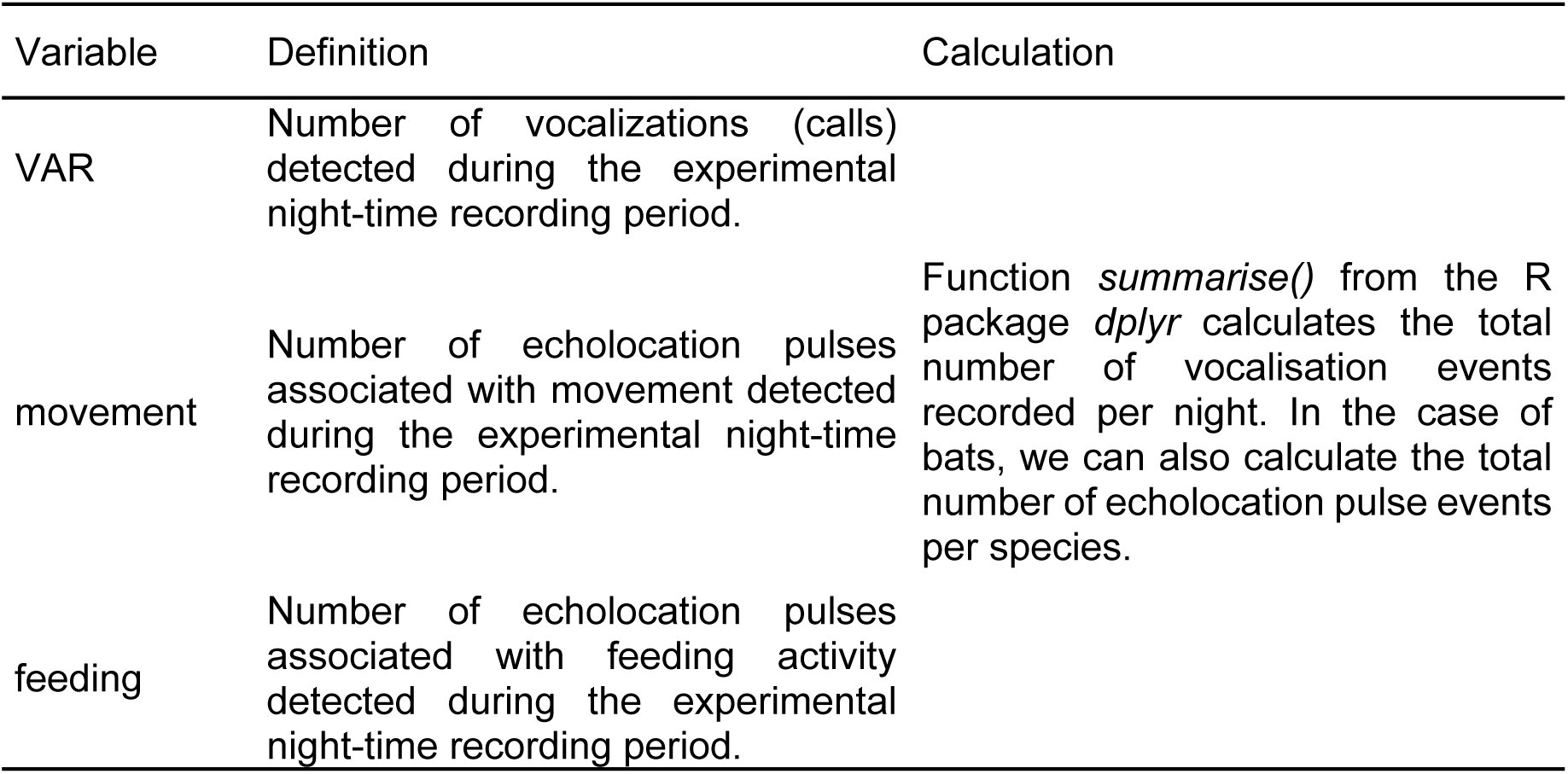
Description of the response variables derived from acoustic recording used for statistical analyses.

All recordings were processed and analysed using Kaleidoscope Pro software (Wildlife Acoustics, version 5.4.9). For Cory’s shearwaters, vocalizations were detected and grouped using the clustering function in Kaleidoscope, which groups acoustically similar calls. Clusters were subsequently inspected by visually examining spectrograms and listening to representative recordings to identify those corresponding to Cory’s shearwater calls, while clusters containing other sounds or noise were discarded.

For bats, ultrasonic recordings were processed using the Kaleidoscope automated species identification (Auto ID) algorithm. All automated identifications were subsequently verified by visually inspecting the spectrograms of each file to confirm species identity and remove false detections, given the known variability in classifier accuracy [50,51]. This procedure ensured that only validated relevant calls associated with the focal species and behaviours were retained for further analysis. In total, 63 nights of Cory’s shearwater recordings (N= 44,936) and 70 nights of bat recordings were included in the analysis (N= 36,814).

### 2.5 Explanatory variables

The main explanatory variable in our study is light treatment, a factor with five levels (i.e. C, AL, AH, WL and WH). Given our field experiment setup, we used five covariates that might influence the individual’s behaviour. These covariates cannot be controlled by the researchers and therefore need to be included in the models (Table 4): 1) Moonlight intensity (moon; lx) modulates the risk of disorientation and attraction to artificial light sources [52], and other general activity patterns such as vocal activity at colony or foraging at sea [38,53]. Likewise, in bats, moonlight intensity can affect foraging activity both directly and indirectly by changing nocturnal insect abundance and behaviour [28,54]. However, artificial light at night can disrupt this natural synchronization, attenuating or even eliminating lunar-driven variations in activity, and thus altering their lunar chronobiology [55]. 2) Temperature (temp; °C), may also play a role in the activity of the study species. In shearwaters, warmer nights might reduce social and vocal activity and even attendance to the colony (authors’ personal observation). In bats, by contrast, warmer nights are known to increase activity due to higher insect availability near lights [56]. 3) Julian date (Jdate) can capture seasonal effects on behaviour during the progression of the annual calendar and the phenology of the species. 4) Year (a two-levels factor) helps to account for inter-annual variation in non-measured variables and ensures that observed effects are not due to annual variability. Finally, 5) location of the two automatic recorders only used for the vocal activity rate of Cory’s shearwaters was included as two-level factor (close and far from experimental lights) to control for potential interactions with treatment.

**Table 4.** Description of the explanatory variables used for statistical models.

| Variable | Definition | Calculation |
| --- | --- | --- |
| moon | A measure of moonlight intensity on the ground (in lux), based on its visible size, the angle above the horizontal and the amount of light it reflects. | Function *calculateMoonlightIntensity()* from R package *moonlit* predicts moonlight intensity on the ground for any given place and time [49]. |
| temp | Physical quantity that expresses the degree of heat or cold of an object or environment (in degrees Celsius). | Requested at State Meteorological Agency (AEMET for its acronym in Spanish), Spanish Government (https://www.aemet.es/es/datos_abiertos/AEMET_OpenData). Temperatures per hour in August 2023 and 2024. |
| Jdate | Julian date. Numerical value from 1 to 365. It allows for a continuous representation of time throughout the year. | Function *yday()* from R package *lubridate* extracts the day of the year from a date object and it assigns a number. |
| year | Year in which the experiment took place. | - |
| location | Two recording devices were used in two locations: one close to the experimental lights and another farther away at a distance of 300 m. |  |

### 2.6 Statistical analyses

Statistical analyses were conducted in *R* version 4.2.3 (R Core Team, 2023). Prior to run the models (see below), all numerical variables were standardized for better comparison of effects sizes (mean = 0, sd = 1). To model the effects of the light treatments on the behaviour of the breeding adult shearwaters, we used *glmmTMB()* function from the *glmmTMB* package [57], for building Generalised Linear Mixed Models (GLMM) including individuals (ID) as a random factor. Our foundational model was y ~ treatment + temperature + year + moon + Jdate + (1|ID). We tested the effect of the treatment, and covariates to control for potential confounding effects. For NV, VAR, movement, and feeding response variables, since they are grouped by night, the random term ID was removed (note that breeders visit the colony no more than once per night), and for VAR specifically, the interaction of light treatment and the location of the automatic recorder was included (treatment*location). Models were fitted using appropriate error distributions and link functions (see <u>Table S2</u>) from *MASS* (version 7.3.58.2) and *stats* (version 4.2.3) R packages [58]. We used the *confint()* function to retrieve 95% confidence intervals (CIs) around the posterior mean estimates and considered support for an effect as strong if zero was not included within the 95% CI.

During model diagnostics, we checked for residual normality and homoscedasticity with *DHARMa* package, version 0.4.7 [59], as well as for multicollinearity issues using the *vif()* function of *car* package, version 3.1-3 [60]. Model diagnosis showed no correlation between any variables, but there is poor residual fit in the movement models of *H. savii* and *P. maderensis* caused by the high variation between years. Post-hoc comparisons among treatments were performed on estimated marginal means (*emmeans*) using Tukey adjustment for multiple comparisons.

## 3. RESULTS

### 3.1 Flight behaviour

We analysed 211 inward flights from 26 Cory’s shearwater breeders. Significant effects of certain light treatments were detected exclusively in TSS, and Altitude models (<u>Table S3</u>; Figure 2). However, post-hoc comparisons failed to detect significant differences among treatments (P-values > 0.1437). Furthermore, for the speed, time, TSS, NDL and NV models, certain secondary variables had a notable influence. In the general models’ predictors, moon (specifically NV), temperature (in time and TSS models) and Julian date (in the case of speed, time, TSS, and NDL) were maintained due to their biological relevance.

**Figure 2.**
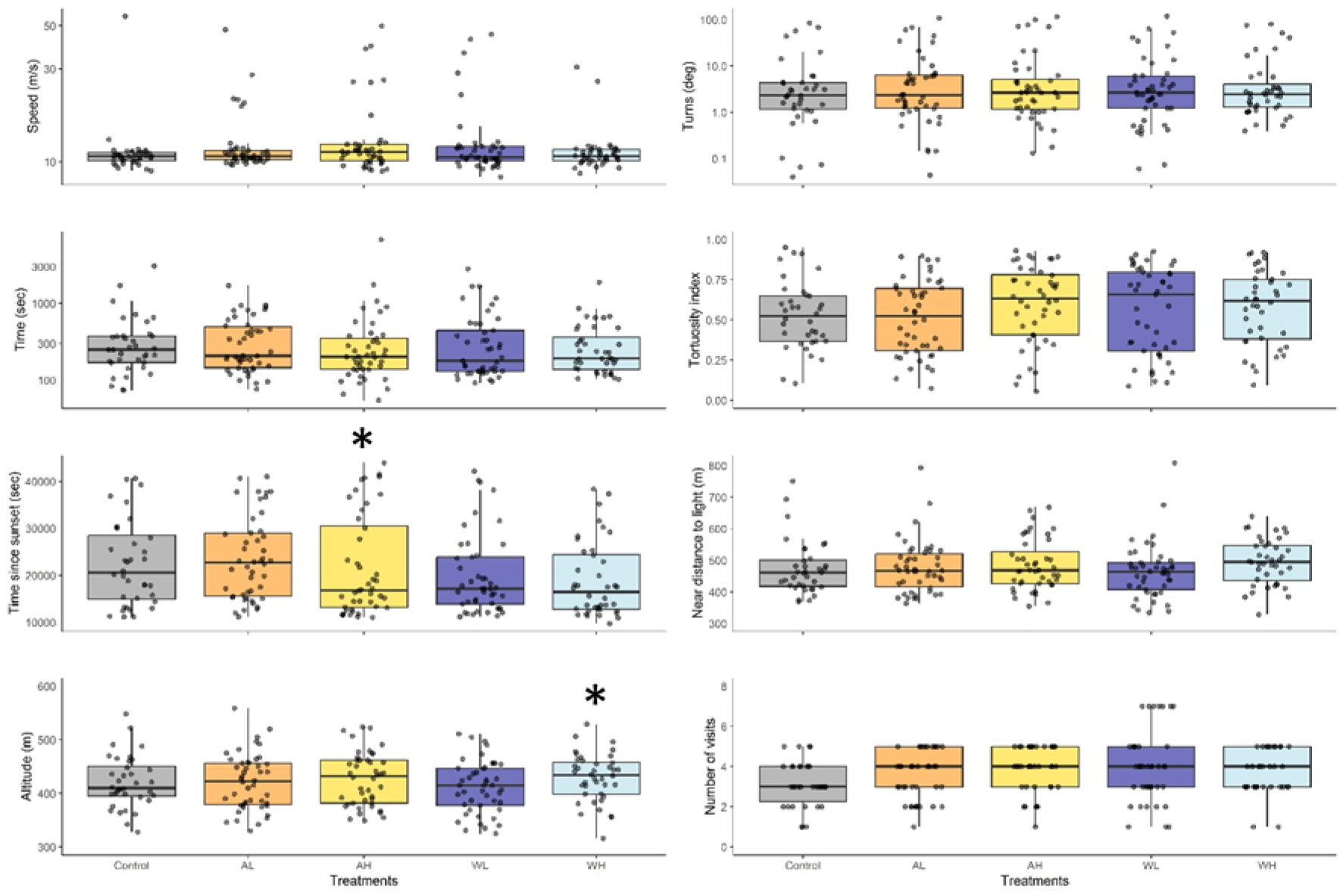
Representation of the effects of light treatments on Cory’s shearwater flight descriptors. Each response variable derived from GPS data is shown along the X-axis, and light treatments along the Y-axis. The response variables include Speed, turning angle (Turns), time spend to enter the colony (time), tortuosity index, time since sunset (TSS), near distance to light (NDL), Altitude and number of visits (NV). Light treatments are: No light (Control), amber low (AL), amber high (AH), white low (WL), white high (WH). Boxplots show medians, quartiles, and 95% confidence intervals; grey-filled dots represent individual data points.

### 3.2 Vocal activity rates

We recorded between 8 and 660 vocalisations of Cory’s shearwater per night (total number = 29251; mean = 240, SD = 130). The impact of experimental light on the rate of vocal activity of shearwaters, during inward flight to the colony, appears to be inconsequential (<u>Table S4</u>; Figure 3). Interestingly, no interaction between light treatment and location of recorders were observed. However, several of the covariates incorporated within the model reached statistical significance, most notably the year of sampling, the Julian date, and the location of the automatic recorder recording higher VAR close to experimental lights. In addition, we tested vocal activity rates of shearwaters combining levels of the treatments of intensity (i.e., high intensity including AH and WH, low intensity including AL and WL, and control) and light colour (i.e., amber including AH and AH, white including WL and WH and control) in two-separated GLMs. No significant differences were observed among these combined treatment levels (results no showed).

**Figure 3.**
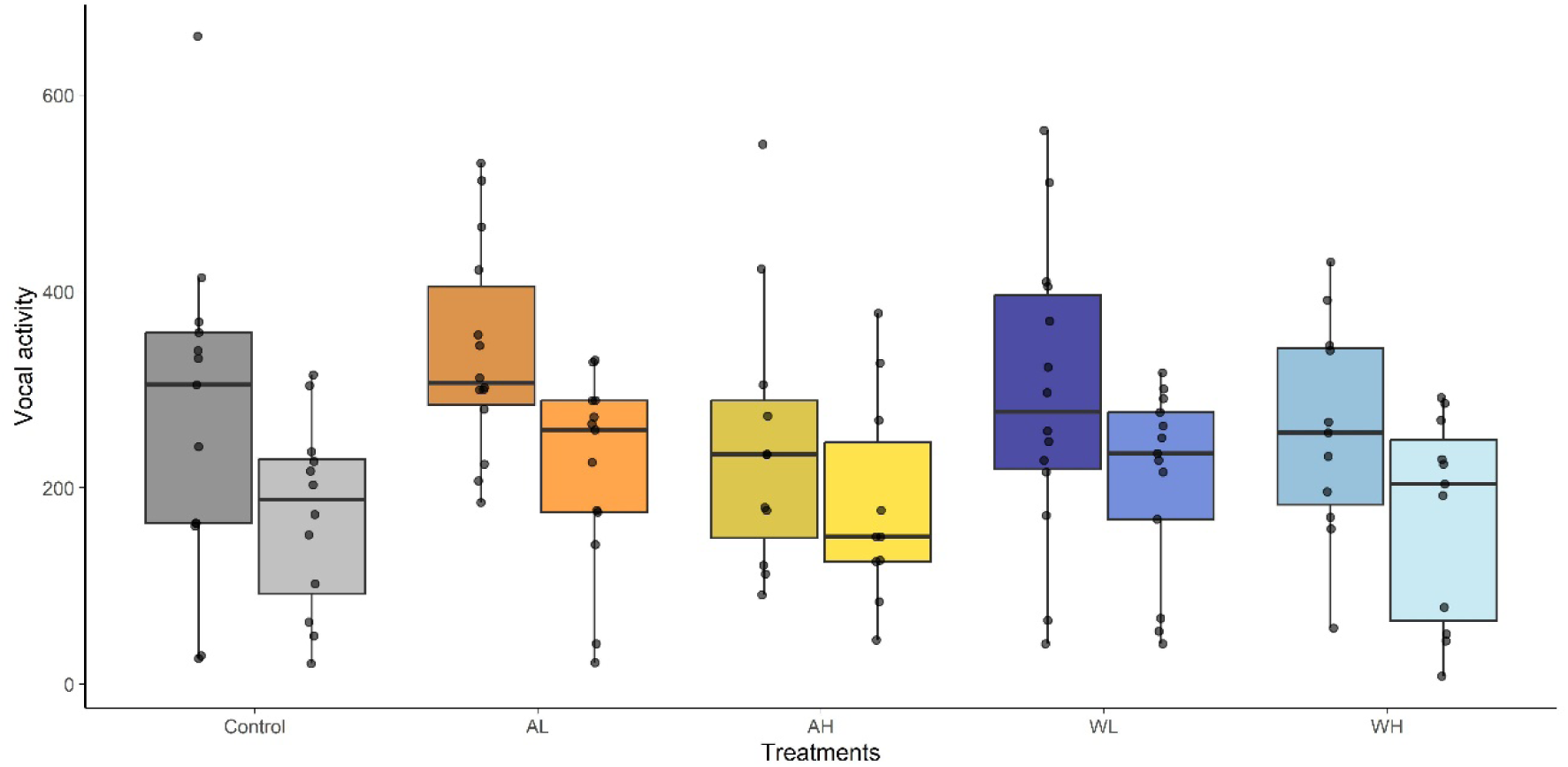
Representation of the vocal activity rate (number of calls recorded during the experimental night-time sampling period; VAR) of Cory’s shearwaters (*Calonectris borealis*) across different light treatments: No light (Control), amber low (AL), amber high (AH), white low (WL), white high (WH). Boxplots show medians, quartiles, and 95% confidence intervals; grey-filled dots represent individual data points. Shading intensity indicates recorder location, with darker tones corresponding to the recorder located close to the experimental lights and lighter tones to the recorder far away from the experimental lights.

Ultrasonic recorders indicated that the study area is inhabited by Savi’s pipistrelle (*Hypsugo savii*; Bonaparte, 1837), Madeiran pipistrelle (*Pipistrellus maderensis*; Dobson, 1878), European free-tailed bat (*Tadarida teniotis*; Rafinesque, 1814), Barbastelle bat (*Barbastella barbastellus*; Schreber, 1774), and Leisler’s bat (*Nyctalus leisleri*; Kuhl, 1817). The abundance of bat recordings was characterized by movement echolocation calls mainly of *H. savii* (<u>Table S5</u>). The abundance of recording of the latter two species was so low that it precluded any statistical analysis and were therefore excluded (<u>Table S5</u>).

Experimental artificial light treatments did not have significant effects on bat activity across most species and behaviour (movement or feeding) analysed. In the feeding and movement models for *H. savii* and *P. maderensis,* the most abundant species according to their detection rates, none of the treatment levels showed statistically significant effects on ultrasonic pulses, suggesting that light treatments did not substantially influence overall bat activity (<u>Table S4</u>; Figure 4). In contrast, in the movement model for *T. teniotis*, low-intensity treatments had a positive effect (although not significant in both cases; see 95% CIs in <u>Table S4</u>), while high-intensity treatments showed marginally negative effects, indicating that low- and high-intensity light treatments may influence the movement in opposite ways in this species. With regards to covariates, year was consistently significant across all models with specific differences effects for the species. Acoustic recordings of *H. savii* and *T. teniotis* were more frequent on the second year (2024) than in the first (2023), while the opposite was true for *P. maderensis* recordings. Julian date was significant in both models for *P. maderensis*, resulting in a decrease in activity as summer progressed. Moon phase showed significant effects in the feeding model of *P. maderensis* (decreasing feeding activity with moonlight) and movement model of *T. teniotis* (increasing activity with moonlight).

**Figure 4.**
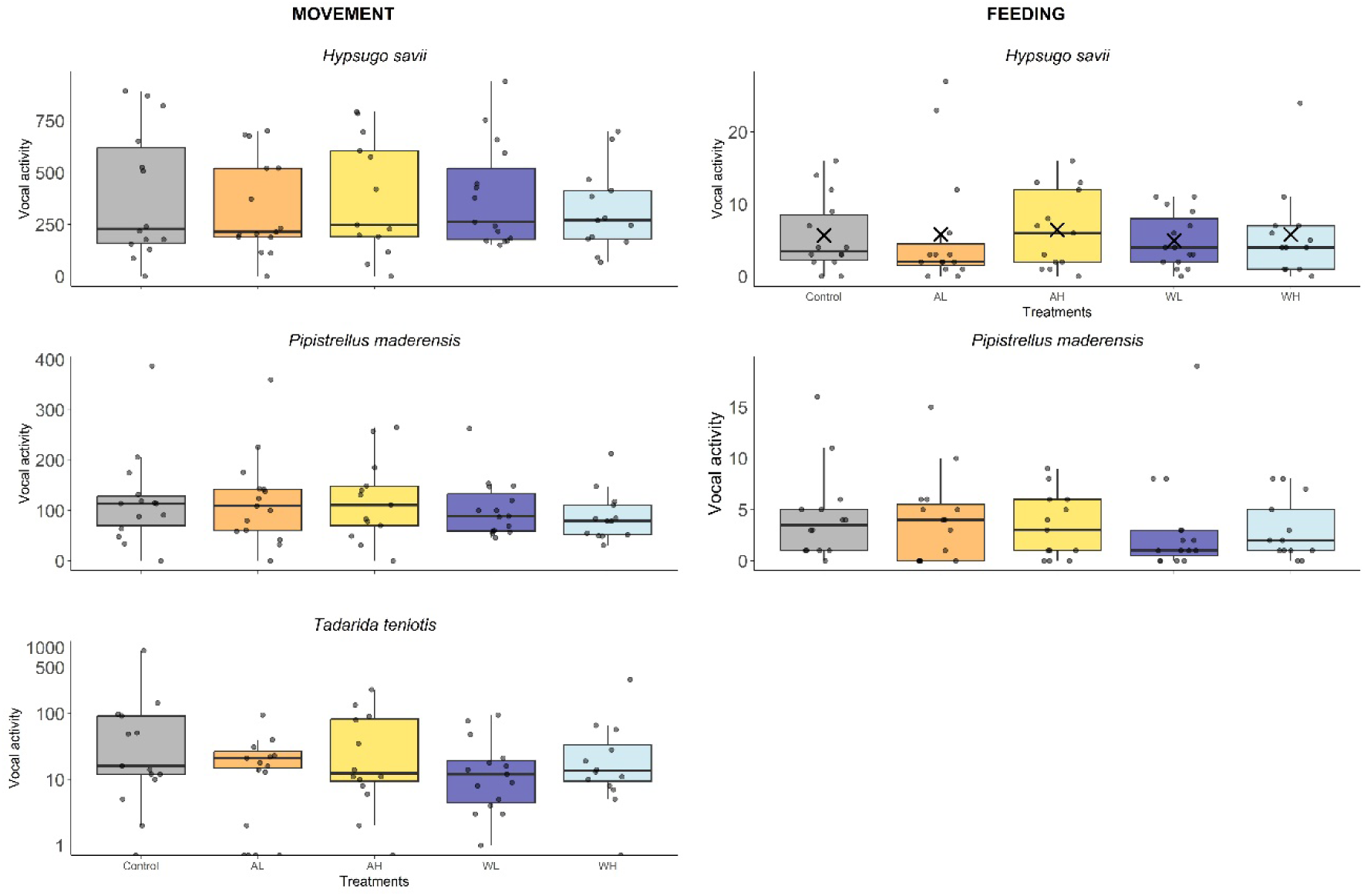
Response of bat species to light treatments. Two behavioural response variables are shown: movement activity and feeding activity, across three bat species: *Pipistrellus maderensis, Hypsugo savii,* and *Tadarida teniotis.* Light treatments are represented on the X-axis as no light (Control), amber low intensity (AL), amber high intensity (AH), white low intensity (WL), and white high intensity (WH). Boxplots display medians, quartiles, and 95% confidence intervals; grey-filled dots represent nightly activity recordings (21 to 00h) for each species.

## 4. DISCUSSION

Despite our efforts to reveal effects of artificial light on seabird and bat activity, we failed to detect significant effects of light treatments in comparison to control (no experimental light) as we predicted. Only a few exceptions reached statistical significance with lack a clear biological interpretation according to the literature background (see hypotheses in the Introduction). In addition, we also failed to detect significant effects among the treatments with the most extreme light combinations allowed under the Canarian Sky Law—those considered potentially the most and least harmful based on the spectrum and intensity of light, i.e., PC amber-low intensity (AL) *vs.* white-high intensity (WH).

Only two out of eight variables describing the flight behaviour of Cory’s shearwaters showed differences among treatments. However, the differences were not in the direction we predicted and are likely spurious results consequence of multiple comparisons. Although the effects were weak and inconsistent, their occurrence under white light broadly aligns with previous experimental studies on adult flying seabirds [23,24,61], where white light treatments have more profound effects than other lights or control treatments.

The differences in the spectral properties of the experimental lights used in our study may not be sufficient to elicit a behavioural response from our animal models, and thus, partly explain these discrepancies. Behavioural responses have been reported under more spectrally contrasting treatments, such as red versus blue light, with adult Cory’s shearwaters showing a preference for red-lit areas [62]. Spectral effects have also been documented in shearwater chicks and fledglings [62,63] and in other seabirds, including penguins and puffins [64,65], suggesting that wavelength composition can play a key role in shaping behavioural responses to artificial light.

Another possible (non-mutually exclusive) explanation is that adult shearwaters may have become accustomed to the light pollution in the surrounding area. The south-west of Tenerife is one of the most light-polluted areas, and it is also where the greatest number of fledglings are grounded by light pollution [41]. Given their longevity, breeding adults may have experienced the progressive increase in artificial light associated with tourist development over recent decades [66] and could have become used to elevated artificial light levels.

In addition to flight behaviour, the rate of vocalizations of Cory’s shearwater also remained largely unaffected by artificial light treatments. In this sense, nocturnal procellariiformes have adapted to the predictable lunar cycle over time, decreasing vocal emissions on bright moonlit nights as a response to minimise predation by visual predators [38]. However, we did not observe a significant decrease in vocalization rates with high moonlight levels. This negative finding supports the idea that adult Cory’s shearwaters may have become accustomed to light pollution, losing natural cues as the high artificial light levels of the near urban areas override the moonlight.

To rule out the influence of prior exposure to light pollution, further research in pristine, non-light-polluted environments is necessary. However, breeding colonies are distributed as they are, and it is hard to find accessible colonies with the necessary logistical facilities (e.g. a power supply) for research purposes whose breeding pairs are not exposed to light pollution [67].

Regarding bats, studies addressing the acoustic activity of the Canarian bat community are virtually absent from the published literature. Our automatic recorders provided information on five bat species. However, frequency of acoustic activity of just three species was suitable for statistical modelling, being *H. savii*, *P. maderensis* and *T. teniotis* the most active. No overall effect of light treatment on acoustic activity was observed across most models. Only *T. teniotis* showed a significant increase in vocal activity under the lowest light-intensity treatment (AL) which is hard to interpret biologically (see below). Similarly to shearwaters, bats could be adapted to an artificially lit landscape. The location of the experimental lights is on the edge of an urban area. Therefore, bats that avoid lights could be repelled by nearby urban lights, while bats that exploit light-attracted insects could visit other permanent lights that are already known as foraging places.

Broader studies indicate that the impact of light pollution on bat activity is highly species and context-dependent [34,68]. Our results are consistent with previous studies suggesting that pipistrelle bats do not alter their movement activity in response to artificial light [69]. However, they do tend to forage more intensively near artificial light sources [70], although this pattern was not reflected in our study. Other studies have shown that bat species within the same genus can exhibit very different foraging behaviours affecting niche segregation, some prefer illuminated habitats, while others avoid them [34,71]. This might explain why we detect mostly *H.* savii and *P. maderensis* species, but not Barbastelle bats, for example. In this study, no feeding buzzes were detected for *T. teniotis*, which may be related to its fast, rectilinear commuting and foraging flights that reduce the probability of detecting terminal buzzes with ground-based acoustic detectors. Additionally, *T. teniotis* showed increased vocal activity under low-light conditions and reduced ultrasonic pulses as light levels rose. This indicates that the species may tolerate, or even prefer, relatively low light levels during flight. This could be due to a trade-off between a lower perceived predation risk or disturbance and the exploitation of light-attracted insects. Similar patterns have been observed in other fast-flying, aerial-hawking bats such as *Nyctalus* and *Eptesicus* spp., which share ecological and morphological traits with *T. teniotis* [72,73]. Our findings suggest that *T. teniotis* adjusts its acoustic activity according to ambient light, with higher vocalization rates when illumination is low and with larger wavelength.

Environmental covariates emerged as significant predictors of bat activity, as predicted. We observed that overall activity declined as Julian date increased, indicating a general seasonal decrease in bat presence or detectability, likely reflecting seasonal shifts in behaviour or activity [74]. In our study, however, we detected no significant effects between temperature and bat activity. This contrasts with other studies, where *P. pipistrellus* has been shown to increase its activity under warmer conditions [56]. These differences may be explained by the Canary Islands’ milder temperatures compared to continental climates in mainland Europe. Finally, moonlight had a significant effect in our models, influencing feeding activity in *P. maderensis* and movement-related ultrasonic pulses in *T. teniotis*. These results suggest that lunar illumination can modulate bat behaviour, although the response may be species and context-specific. Previous studies have reported mixed findings, with some documenting behavioural effects of moonlight and others finding no evidence of lunar phobia in swarming bats [28,75].

To our best knowledge, our study constitutes one of the first experimental efforts to test the adequacy of astronomer-led artificial light regulations on biodiversity. We failed to detect substantial effects of the most extreme light combinations allowed under the Canarian Sky Law. In addition, we contribute empirical data on nocturnal wildlife (GPS tracks of shearwaters and recordings of acoustic activity of a bat community) on light polluted nightscapes. Future studies with extended monitoring periods to increase sample size and developed in more pristine study areas to involve animals non previously exposed to light pollution are essential to clarify the subtle and potentially species-specific impacts of artificial light regulations on nocturnal wildlife.

## 5. ACKNOWLEGMENTS

We are grateful to many friends to helping during fieldwork: Ingrid Rivero, Ruth Martínez, Miguel Velázquez, Rayco Velázquez, Pablo J. Vázquez, Juan Curbelo, Enrique Sacramento and Abraham Hernández. We are grateful to Ayuntamiento de Adeje, Restaurante Otelo, and Andrei (Barranco del Infierno) for helping us with logistics. Salva Bará provided useful comments on a draft of this manuscript.

## Author contributions

Conceptualization: A.R., B.R.; Methodology: A.R, B.R., D.G., C.dT., J.F.dlP.; Fieldwork: B.R., A.R.; Formal analysis: C.dT., D.G.; Resources: A.R., D.G.; Data curation: C.dT., D.G.; Writing - original draft: C.dT., A.R.; Writing - review & editing: A.R., B.R., D.G., J.F.dlP.; Supervision: A.R.; Project administration: A.R.; Funding acquisition: A.R.

## Funding

This study was supported by the project LightingBirds PID2021-124101OA-I00, funded by MCIN/AEI/10.13039/501100011033/ FEDER, UE, granted to AR. C.d.T was supported by a JAE intro CSIC fellowship (JAEINT_23_02175). A.R. was supported by the European Union under the Horizon Europe Programme, grant agreement no. 101135471 (AquaPLAN) and contracted by a Ramón y Cajal fellowship (RYC2021-032656-I) funded by Ministerio de Ciencia, Innovación y Universidades MICIU/AEI/10.13039/501100011033 and the European Union «NextGenerationEU»/PRTR.

## Data availability

GPS data are available at Seabird Tracking Database, BirdLife International; data set ID numbers 2298 and 2299.

Acoustic recording of Cory’s shearwater and bat community are available upon request.

## Competing interests

The authors declare no competing or financial interests.

## Notes

### Competing Interest Statement

The authors have declared no competing interest.

## REFERENCES

1. Riegel KW. Light Pollution. Science. 1973;179: 1285–1291. doi:10.1126/science.179.4080.1285

2. Varela AM. The increasing effects of light pollution on professional and amateur astronomy. Science. 2023;380: 1136–1140. doi:10.1126/science.adg0269

3. Barentine JC. Who speaks for the night? The regulation of light pollution in the ‘Rights of Nature’ legal framework. Int J Sustain Light. 2020;22: 28–36. doi:10.26607/ijsl.v22i2.104

4. Falchi F, Cinzano P, Elvidge CD, Keith DM, Haim A. Limiting the impact of light pollution on human health, environment and stellar visibility. J Environ Manage. 2011;92: 2714–2722. doi:10.1016/j.jenvman.2011.06.029

5. Falchi F, Bará S. Light pollution is skyrocketing. Science. 2023;379: 234–235. doi:10.1126/science.adf4952

6. Kyba CCM, Kuester T, Sánchez de Miguel A, Baugh K, Jechow A, Hölker F, et al. Artificially lit surface of Earth at night increasing in radiance and extent. Sci Adv. 2017;3: e1701528. doi:10.1126/sciadv.1701528

7. Kyba CCM, Altıntaş YÖ, Walker CE, Newhouse M. Citizen scientists report global rapid reductions in the visibility of stars from 2011 to 2022. Science. 2023;379: 265–268. doi:10.1126/science.abq7781

8. Jones TM, McNamara KB. Ecological light pollution. Curr Biol. 2023;33: R843–R844. doi:10.1016/j.cub.2023.06.034

9. Gaston KJ, Visser ME, Hölker F. The biological impacts of artificial light at night: the research challenge. Philos Trans R Soc B Biol Sci. 2015;370: 20140133. doi:10.1098/rstb.2014.0133

10. Longcore T, Rich C. Ecological light pollution. Front Ecol Environ. 2004;2: 191–198. doi:10.1890/1540-9295(2004)002[0191:ELP]2.0.CO;2

11. Hölker F, Wolter C, Perkin EK, Tockner K. Light pollution as a biodiversity threat. Trends Ecol Evol. 2010;25: 681–682. doi:10.1016/j.tree.2010.09.007

12. Gaston KJ, Sánchez de Miguel A. Environmental Impacts of Artificial Light at Night. Annu Rev Environ Resour. 2022;47: 373–398. doi:10.1146/annurev-environ-112420-014438

13. Hölker F, Jechow A, Schroer S, Tockner K, Gessner MO. Light pollution of freshwater ecosystems: principles, ecological impacts and remedies. Philos Trans R Soc B Biol Sci. 2023;378. doi:10.1098/rstb.2022.0360

14. Marangoni LFB, Davies T, Smyth T, Rodríguez A, Hamann M, Duarte C, et al. Impacts of artificial light at night in marine ecosystems—A review. Glob Change Biol. 2022;28: 5346–5367. doi:10.1111/gcb.16264

15. Hirt MR, Evans DM, Miller CR, Ryser R. Light pollution in complex ecological systems. Philos Trans R Soc B Biol Sci. 2023;378: 20220351. doi:10.1098/rstb.2022.0351

16. Gaston KJ, Davies TW, Bennie J, Hopkins J. Review: Reducing the ecological consequences of night-time light pollution: options and developments. J Appl Ecol. 2012;49: 1256–1266. doi:10.1111/j.1365-2664.2012.02212.x

17. Hölker F, Moss T, Griefahn B, Kloas W, Voigt CC, Henckel D, et al. The Dark Side of Light : A Transdisciplinary Research Agenda for Light. Ecol Soc. 2010;15: 13. doi:10.1890/080129

18. Benzaken D (Dominique), Renard I. Future directions for biodiversity action in Europe overseas: outcomes of the Review of the Implementation of the Convention on Biological Diversity, December 2010. Gland, Switzerland: IUCN; 2011. Available: https://iucn.org/resources/publication/future-directions-biodiversity-action-europe-overseas-outcomes-review

19. Imber MJ. Behaviour of petrels in relation to the Moon and artificial lights. Notornis, Journal of the Ornithological Society of New Zealand. 1975;22: 302–306.

20. Rodríguez A, Holmes ND, Ryan PG, Wilson K-J, Faulquier L, Murillo Y, et al. Seabird mortality induced by land-based artificial lights. Conserv Biol. 2017;31: 986–1001. doi:10.1111/cobi.12900

21. Dick MH, Donaldson W. Fishing Vessel Endangered by Crested Auklet Landings. The Condor. 1978;80: 235. doi:10.2307/1367924

22. Raine AF, Driskill S, Rothe J, Rossiter S, Gregg J, Anderson T, et al. The Impact of Light Attraction on Adult Seabirds and the Effectiveness of Minimization Actions. Pac Sci. 2024;78: 85–102. doi:10.2984/78.1.6

23. Middlemiss KL, Cieraad E, Mander S, Fischer JH, Goad D. Understanding the impacts of LED light pollution in marine ecosystems: phototaxis response in fairy prion. J Ornithol. 2025;166: 677–689. doi:10.1007/s10336-024-02245-1

24. Austad M, Oppel S, Crymble J, Greetham HR, Sahin D, Lago P, et al. The effects of temporally distinct light pollution from ships on nocturnal colony attendance in a threatened seabird. J Ornithol. 2023;164: 527–536. doi:10.1007/s10336-023-02045-z

25. Cianchetti-Benedetti M, Becciu P, Massa B, Dell’Omo G. Conflicts between touristic recreational activities and breeding shearwaters: short-term effect of artificial light and sound on chick weight. Eur J Wildl Res. 2018;64: 19. doi:10.1007/s10344-018-1178-x

26. Polak T, Korine C, Yair S, Holderied MW. Differential effects of artificial lighting on flight and foraging behaviour of two sympatric bat species in a desert. J Zool. 2011;285: 21–27. doi:10.1111/j.1469-7998.2011.00808.x

27. Gili F, Fassone C, Rolando A, Bertolino S. In the Spotlight: Bat Activity Shifts in Response to Intense Lighting of a Large Railway Construction Site. Sustainability. 2024;16: 2337. doi:10.3390/su16062337

28. Saldaña-Vázquez RA, Munguía-Rosas MA. Lunar phobia in bats and its ecological correlates: A meta-analysis. Mamm Biol. 2013;78: 216–219. doi:10.1016/j.mambio.2012.08.004

29. Pauwels J, Le Viol I, Bas Y, Valet N, Kerbiriou C. Adapting street lighting to limit light pollution’s impacts on bats. Glob Ecol Conserv. 2021;28: e01648. doi:10.1016/j.gecco.2021.e01648

30. Hale JD, Fairbrass AJ, Matthews TJ, Davies G, Sadler JP. The ecological impact of city lighting scenarios: exploring gap crossing thresholds for urban bats. Glob Change Biol. 2015;21: 2467–2478. doi:10.1111/gcb.12884

31. Rydell J, Eklöf J, Sánchez-Navarro S. Age of enlightenment: long-term effects of outdoor aesthetic lights on bats in churches. R Soc Open Sci. 2017;4: 161077. doi:10.1098/rsos.161077

32. Stone EL, Harris S, Jones G. Impacts of artificial lighting on bats: a review of challenges and solutions. Mamm Biol. 2015;80: 213–219. doi:10.1016/j.mambio.2015.02.004

33. Straka TM, Wolf M, Gras P, Buchholz S, Voigt CC. Tree Cover Mediates the Effect of Artificial Light on Urban Bats. Front Ecol Evol. 2019;7. doi:10.3389/fevo.2019.00091

34. Voigt CC, Dekker J, Fritze M, Gazaryan S, Hölker F, Jones G, et al. The Impact of Light Pollution on Bats Varies According to Foraging Guild and Habitat Context. BioScience. 2021;71: 1103–1109. doi:10.1093/biosci/biab087

35. Longcore T, Rodríguez A, Witherington B, Penniman JF, Herf L, Herf M. Rapid assessment of lamp spectrum to quantify ecological effects of light at night. J Exp Zool Part Ecol Integr Physiol. 2018;329: 511–521. doi:10.1002/jez.2184

36. Longcore T. A compendium of photopigment peak sensitivities and visual spectral response curves of terrestrial wildlife to guide design of outdoor nighttime lighting. Basic Appl Ecol. 2023;73: 40–50. doi:10.1016/j.baae.2023.09.002

37. Davies TW, Bennie J, Inger R, De Ibarra NH, Gaston KJ. Artificial light pollution: are shifting spectral signatures changing the balance of species interactions? Glob Change Biol. 2013;19: 1417–1423. doi:10.1111/gcb.12166

38. Bretagnolle V. Effet de la lune sur l’activité des pétrels (classe Aves) aux îles Salvages (Portugal). Can J Zool. 1990;68: 1404–1409. doi:10.1139/z90-209

39. Instituto de Estadística de Canarias (ISTAC). Población según sexos y edades. Canarias, islas y municipios por años. In: Banco de datos [Internet]. 2024 [cited 5 June 2025]. Available: https://www3.gobiernodecanarias.org/istac/statistical-visualizer/visualizer/data.html?resourceType=dataset&agencyId=ISTAC&resourceId=E30243A_000001&version=~latest#visualization/table

40. Reyes-González JM, González-Solís J, Salvador Milla A. Pardela cenicienta atlántica – Calonectris borealis (Cory, 1881). 2016 [cited 20 June 2025]. doi:10.20350/digitalCSIC/8918

41. Rodríguez A, Rodríguez B. Attraction of petrels to artificial lights in the Canary Islands: effects of the moon phase and age class. Ibis. 2009;151: 299–310. doi:10.1111/j.1474-919X.2009.00925.x

42. Fontaine R, Gimenez O, Bried J. The impact of introduced predators, light-induced mortality of fledglings and poaching on the dynamics of the Cory’s shearwater (*Calonectris diomedea*) population from the Azores, northeastern subtropical Atlantic. Biol Conserv. 2011;144: 1998–2011. doi:10.1016/j.biocon.2011.04.022

43. Rodríguez A, García D, Rodríguez B, Cardona E, Parpal L, Pons P. Artificial lights and seabirds: is light pollution a threat for the threatened Balearic petrels? J Ornithol. 2015;156: 893–902. doi:10.1007/s10336-015-1232-3

44. Ramírez-Fráncel LA, García-Herrera LV, Losada-Prado S, Reinoso-Flórez G, Sánchez-Hernández A, Estrada-Villegas S, et al. Bats and their vital ecosystem services: a global review. Integr Zool. 2022;17: 2–23. doi:10.1111/1749-4877.12552

45. Tuneu-Corral C, Puig-Montserrat X, Riba-Bertolín D, Russo D, Rebelo H, Cabeza M, et al. Pest suppression by bats and management strategies to favour it: a global review. Biol Rev. 2023;98: 1564–1582. doi:10.1111/brv.12967

46. Fajardo S, Benzal Pérez J. Datos sobre la distribución de quirópteros en Canarias (Mammalia: Chiroptera). Vieraea Folia Sci Biol Canar. 2002; 213–230.

47. Russo D, Cistrone L. IUCN Red List of Threatened Species: Plecotus teneriffae. IUCN Red List Threat Species. 2023 [cited 14 Nov 2025]. Available: https://www.iucnredlist.org/en

48. Wilson R, Pütz K, Peters G, Culik B, Scolaro A, Charrassin J-B, et al. Long term attachment of transmitting and recording devices to penguins and other seabirds. Wildlife Society Bulletin. 1 Mar 1997;25: 101–106.

49. Śmielak MK. Biologically meaningful moonlight measures and their application in ecological research. Behav Ecol Sociobiol. 2023;77: 21. doi:10.1007/s00265-022-03287-2

50. Roby PL, Jordan GW. Importance of Manually Vetting Acoustic Bat Call Files: A Case Study for Northern Long-Eared Bats. J Fish Wildl Manag. 2024;15: 510–518. doi:10.3996/JFWM-23-046

51. Solick DI, Hopp BH, Chenger J, Newman CM. Automated echolocation classifiers vary in accuracy for northeastern U.S. bat species. PLOS ONE. 2024;19: e0300664. doi:10.1371/journal.pone.0300664

52. Rodríguez A, Atchoi E, Rodríguez B, Pipa T, Le Corre M, Ainley DG. Moonlight diminishes seabird attraction to artificial light. Conserv Sci Pract. 2023;5: e13014. doi:10.1111/csp2.13014

53. Rubolini D, Maggini I, Ambrosini R, Imperio S, Paiva VH, Gaibani G, et al. The Effect of Moonlight on Scopoli’s Shearwater Calonectris diomedea Colony Attendance Patterns and Nocturnal Foraging: A Test of the Foraging Efficiency Hypothesis. Ethology. 2015;121: 284–299. doi:10.1111/eth.12338

54. Appel G, López-Baucells A, Ernest-Magnusson W, Bobrowiec PED. Aerial insectivorous bat activity in relation to moonlight intensity. Mamm Biol. 2017;85: 37–46. doi:10.1016/j.mambio.2016.11.005

55. Li H, Allen P, Boris S, Lagrama S, Lyons J, Mills C, et al. Artificial light at night (ALAN) pollution alters bat lunar chronobiology: insights from broad-scale long-term acoustic monitoring. Ecol Process. 2024;13: 13. doi:10.1186/s13717-024-00491-y

56. Buddendorf SWJ, Visser ME, Spoelstra K. Temperature amplifies the effect of anthropogenic light on foraging common pipistrelle bats. Biol Lett. 2025;21: 20250049. doi:10.1098/rsbl.2025.0049

57. Brooks M, Bolker B, Kristensen K, Maechler M, Magnusson A, McGillycuddy M, et al. glmmTMB: Generalized Linear Mixed Models using Template Model Builder. 2025. Available: https://cran.r-project.org/web/packages/glmmTMB/index.html

58. Venables WN, Ripley BD. Modern applied statistics with S. New York: Springer; 2002. Available: https://cran.r-project.org/web/packages/MASS/citation.html

59. Hartig F, Lohse L, leite M de S. DHARMa: Residual Diagnostics for Hierarchical (Multi-Level / Mixed) Regression Models. 2024. Available: https://cran.r-project.org/web/packages/DHARMa/index.html

60. Fox J, Weisberg S. An R Companion to Applied Regression. 3rd ed. Thousand Oaks, CA: Sage; 2019.

61. Syposz M, Padget O, Willis J, Van Doren BM, Gillies N, Fayet AL, et al. Avoidance of different durations, colours and intensities of artificial light by adult seabirds. Sci Rep. 2021;11: 18941. doi:10.1038/s41598-021-97986-x

62. Atchoi E, Mitkus M, Machado B, Medeiros V, Garcia S, Juliano M, et al. Do seabirds dream of artificial lights? Understanding light preferences of Procellariiformes. J Exp Biol. 2024;227. doi:10.1242/jeb.247665

63. Atchoi E, Mitkus M, Vitta P, Machado B, Rocha M, Juliano M, et al. Ontogenetic exposure to light influences seabird vulnerability to light pollution. J Exp Biol. 2023;226: jeb245126. doi:10.1242/jeb.245126

64. Brown TM, Wilhelm SI, Slepkov AD, Baker K, Mastromonaco GF, Burness G. Navigating the night: effects of artificial light on the behaviour of Atlantic puffin fledglings. Anim Behav. 2024;218: 135–148. doi:10.1016/j.anbehav.2024.09.008

65. Rodríguez A, Holmberg R, Dann P, Chiaradia A. Penguin colony attendance under artificial lights for ecotourism. J Exp Zool Part Ecol Integr Physiol. 2018;329. doi:10.1002/jez.2155

66. Rodríguez A, Rodríguez B, Lucas MP. Trends in numbers of petrels attracted to artificial lights suggest population declines in Tenerife, Canary Islands. Ibis. 2012;154: 167–172. doi:10.1111/j.1474-919X.2011.01175.x

67. Reyes-González JM, González-Solís J. Pardela cenicienta atlántica – *Calonectris borealis*. In: Salvador A, Morales MB, editors. Enciclopedia Virtual de los Vertebrados Españoles. Madrid: Museo Nacional de Ciencias Naturales; 2016.

68. Barré K, Vernet A, Azam C, Le Viol I, Dumont A, Deana T, et al. Landscape composition drives the impacts of artificial light at night on insectivorous bats. Environ Pollut. 2022;292: 118394. doi:10.1016/j.envpol.2021.118394

69. Russo D, Cosentino F, Festa F, De Benedetta F, Pejic B, Cerretti P, et al. Artificial illumination near rivers may alter bat-insect trophic interactions. Environ Pollut. 2019;252: 1671–1677. doi:10.1016/j.envpol.2019.06.105

70. Voigt CC, Scholl JM, Bauer J, Teige T, Yovel Y, Kramer-Schadt S, et al. Movement responses of common noctule bats to the illuminated urban landscape. Landsc Ecol. 2020;35: 189–201. doi:10.1007/s10980-019-00942-4

71. Salinas-Ramos VB, Ancillotto L, Cistrone L, Nastasi C, Bosso L, Smeraldo S, et al. Artificial illumination influences niche segregation in bats. Environ Pollut. 2021;284: 117187. doi:10.1016/j.envpol.2021.117187

72. Hermans C, Kijm L, Paardekooper M, Koblitz JC, Stilz P, Haarsma A-J, et al. Limited immediate effect of artificial light of realistic intensity on flight behaviour of commuting pond bat (*Myotis dasycneme*). Basic Appl Ecol. 2025;87: 20–28. doi:10.1016/j.baae.2025.05.007

73. Hooker J, Lintott P, Stone E. Lighting up our waterways: Impacts of a current mitigation strategy on riparian bats. Environ Pollut. 2022;307: 119552. doi:10.1016/j.envpol.2022.119552

74. Silva MC, Granadeiro JP, Boersma PD, Strange I. Effects of predation risk on the nocturnal activity budgets of thin-billed prions Pachyptila belcheri on New Island, Falkland Islands. Polar Biol. 2011;34: 421–429. doi:10.1007/s00300-010-0897-6

75. Apoznański G, Tuff F, Carr A, Rachwald A, Marszałek E, Marszałek T, et al. Absence of lunar phobia in European swarming vespertilionid bats. Sci Rep. 2024;14: 2675. doi:10.1038/s41598-024-53281-z

